# doubleHelix: nucleic acid sequence identification, assignment and validation tool for cryo-EM and crystal structure models

**DOI:** 10.1101/2023.02.17.528963

**Authors:** Grzegorz Chojnowski

**Affiliations:** European Molecular Biology Laboratory, c/o DESY, Notkestraße 85, 22607 Hamburg, Germany

**Keywords:** cryo-EM, MX, nucleic acids, sequence assignment, model validation

## Abstract

Sequence assignment is a key step of the model building process in both cryogenic electron microscopy (cryo-EM) and macromolecular crystallography (MX). If the assignment fails, it can result in difficult to identify errors affecting the interpretation of a model. There are many model validation strategies that help experimentalists in this step of protein model building, but they are virtually non-existent for nucleic acids. Here I present doubleHelix – a comprehensive method for assignment, identification, and validation of nucleic acid sequences in structures determined using cryo-EM and MX. The method combines a neural network classifier of nucleobase identities and a sequence-independent secondary structure assignment approach. I show that the presented method can successfully assist model building at lower resolutions, where visual map interpretation is very difficult. Moreover, I present examples of sequence assignment errors detected using doubleHelix in cryo-EM and MX structures of ribosomes deposited in the Protein Data Bank, which escaped the scrutiny of available model-validation approaches.

The doubleHelix program source code is available under BSD-3 license at https://gitlab.com/gchojnowski/doublehelix.

## INTRODUCTION

Nucleic acids are key players in many cellular processes ranging from gene expression to the catalysis of chemical reactions. For many nucleic acid molecules tertiary structure determines function, much like for proteins. Nevertheless, our understanding of the structure-function relationship in nucleic acids clearly lags behind proteins, which is reflected by the disproportion of structures deposited in the Protein Data Bank (PDB) (1). As of January 2023, out of 200,708 available structures only 15,374 (7%) contained a nucleic acid component. The resolution revolution in cryogenic electron microscopy (cryo-EM) seems to be slowly changing this picture as more and more challenging nucleic-acid complexes are being determined using this technique. In 2022, out of 1,454 structures with nucleic-acid components deposited in the PDB as many as 804 (55%) were determined using cryo-EM. Many of these cryo-EM structures would be very difficult to solve using other techniques owing to their size and structural heterogeneity (2).

The release of the Artificial Intelligence (AI) based structure prediction programs AlphaFold2 (3) and RoseTTAFold (4) provided means for accurate and widely accessible structure prediction of protein structures. Although they did not solve the problem of protein structure determination the accurate predictions proved very effective in assisting experimental structure determination (5). They are robust Molecular Replacement search models for solving the phase problem in macromolecular X-ray crystallography (6). Owing to their reliability AlphaFold2 predictions also greatly facilitate interpretation of cryo-EM maps, in particular of huge complexes like a spectacular example of a recently published structure of Nuclear Pore Complex (7). Moreover, AI-based predictions fitted into cryo-EM and MX maps usually need little rebuilding and refinement (6,8), increasing the speed at which a final model can be obtained.

There is no AlphaFold2 or RoseTTAFold equivalent for nucleic acids and most likely won’t be soon as building AI 3D structure prediction tools requires huge and diverse training sets of reliable structural models that are currently not available for nucleic acids. The PDB-deposited models are strongly biased towards ribosomal RNA that is usually highly conserved across all kingdoms of live. Moreover, as will be shown later in this work, available experimental RNA structures contain difficult to identify errors that may reduce generalization properties of structure prediction methods. Although attempts to build such tools are already taken (9), experimental techniques will continue to be the method of choice for detailed structural studies of nucleic acid complexes, with all their limitations and bottlenecks.

MX and cryo-EM remain the most frequently used experimental approaches for the structure determination of large biomolecules. The main result of both these methods is an atomic model traced into a map - an interpretation of experimental observations given *a priori* knowledge of biomolecular structure. Although, the main effort of the method developer community is clearly focused on proteins, several techniques facilitating experimental determination of nucleic acid structures in cryo-EM and MX have been developed, e.g. NAUTILUS (10), ARP/wARP (11), PHENIX (12), RCrane (13), COOT (14), ISOLDE (15). As with proteins, nucleic acid model building in these methods usually starts with tracing into a density map a ribose-phosphate backbone that makes up two-thirds of a polynucleotide mass. The backbone model is subsequently assigned to a target sequence and complemented with base moieties.

Sequence assignment is a crucial step in macromolecular model building. It is required for the identification and completion of missing fragments in initial models. It is also a fundamental prerequisite of a model interpretation. Failure may lead to register-shift errors, where residues are systematically assigned an identity of a residue a few positions before or ahead in sequence. Although register-shifts may bias model interpretation, they remain one of the most difficult problems to identify and correct in macromolecular models (16). In protein models, register-shifts often result in backbone-geometry outliers when several sidechains are forced into too small density volumes, which can be detected using geometry validation approaches like CaBLAM (17). Moreover, backbone tracing issues (e.g. deletion or insertion) that caused register shift can be occasionally detected as a geometry outlier (18). Nevertheless, it has been shown that regardless of the effort made to validate protein models, register-shift errors are relatively common in PDB (19,20). Particularly affected are very large structures, for example ribosomes, where detailed inspection of a model using interactive tools (e.g. COOT or ISOLDE) is rarely feasible.

Sequence assignment errors in nucleic acids are even more difficult to identify than in proteins. They rarely result in severe geometry issues during model refinement as the ribose-phosphate backbone dominates scattering and its geometry is weakly affected by the presence of misassigned nucleobases. Moreover, small differences between different types of purines and pyrimidines makes visual validation of a sequence assignment very challenging unless high-resolution maps are available.

The most prominent issue related to a sequence assignment error in nucleic acid structures are steric clashes arising from the presence of base-paired nucleobases that don’t fit their secondary structure context - non-isostericity (21). For example, a Watson-Crick pair in cis orientation that erroneously involves two guanines is too large to fit into a double-helical region (22). At lower resolutions, however, this will be promptly masked by a refinement software and the bases that are weakly restrained by a map shifted to a non-clashing conformation. Nevertheless, these relatively rare issues can be in principle be detected using standard model-validation software, e.g. Molprobity (17).

Another, rarely recognised, issue related to the sequence assignment in model building are unknown target sequences. Until recently, structural studies of macromolecules of unknown identity, e.g. extracted from natural sources, were attempted predominantly using MX (23). Recent developments in cryo-EM, however, forged a completely new path to the studies of uncharacterised macromolecules. It has been shown that cryo-EM reconstructions of protein nucleic-acid complexes, at resolutions high enough for *de-novo* model building, can be determined directly in a cell using subtomogram averaging (24). High-resolution structural information can be also retrieved from a systematic cryo-EM analysis of cell lysate fractions (25,26). In a recent study Skalidis and co-workers (27) presented a complete workflow for identifying biomolecules directly from native cell extracts combining cryo-EM with AI structure prediction methods. Although there are in principle no technical limitations to the identification of nucleic-acid sequences directly from cryo-EM reconstruction, to the best of my knowledge there is no computational tool available that could be used for this purpose.

In this work I present *doubleHelix*; a computer program for comprehensive nucleic-acid sequence identification, assignment, and validation in MX and cryo-EM models. Similarly to a previously developed program *findMySequence* (28) for protein-sequence identification, doubleHelix uses neural network classifiers for estimating residue-type probabilities given a backbone model and a density map. What makes the doubleHelix program unique is the way it addresses the inherent nucleobase-type ambiguity that makes it impossible to distinguish adenine from guanine and cytosine from uracil or thymine unless a very high-resolution experimental data is available. The program estimates only the probabilities of purines and pyrimidines in a model. This information is complemented with base-pairing restraints obtained using a new approach that relies on a backbone conformation ignoring nucleobase identities that are not known before the sequence assignment. The base-pair identification approach is based on alignment of recurrent nucleic-acid structural motifs of known secondary structure (e.g. A- or B-form double helices) to the target model. I show that despite its simplicity this approach is both highly specific and accurate. Moreover, the secondary structure information it provides readily improves sequence assignment and identification performance at lower resolutions where base-type classification reliability is reduced. I also show an example of a previously unidentified RNA-sequence assignment errors in mammalian and bacterial ribosome structures deposited in the PDB that could have been avoided if *doubleHelix* had been used for model building and validation.

## AN OVERVIEW OF THE DOUBLEHELIX METHOD

The *doubleHelix* program requires on input a model in PDB or mmCIF format. For the sequence identification and assignment, it also requires a corresponding map, which can be provided in CCP4/MRC format for cryo-EM models or as a MTZ file with structure factor amplitudes and phases for crystal structures. The doubleHelix program provides four basic functionalities:

### Secondary structure restraints generator for nucleic-acid models

Given RNA or DNA model on input the program generates base-pair and -stacking restraints in formats accepted by COOT and popular refinement programs REFMAC5 (29) and PHENIX (30). Additionally, it generates a PYMOL (31) script that can be used for visualising the restraints (Figure 1C).

**Figure 1.**
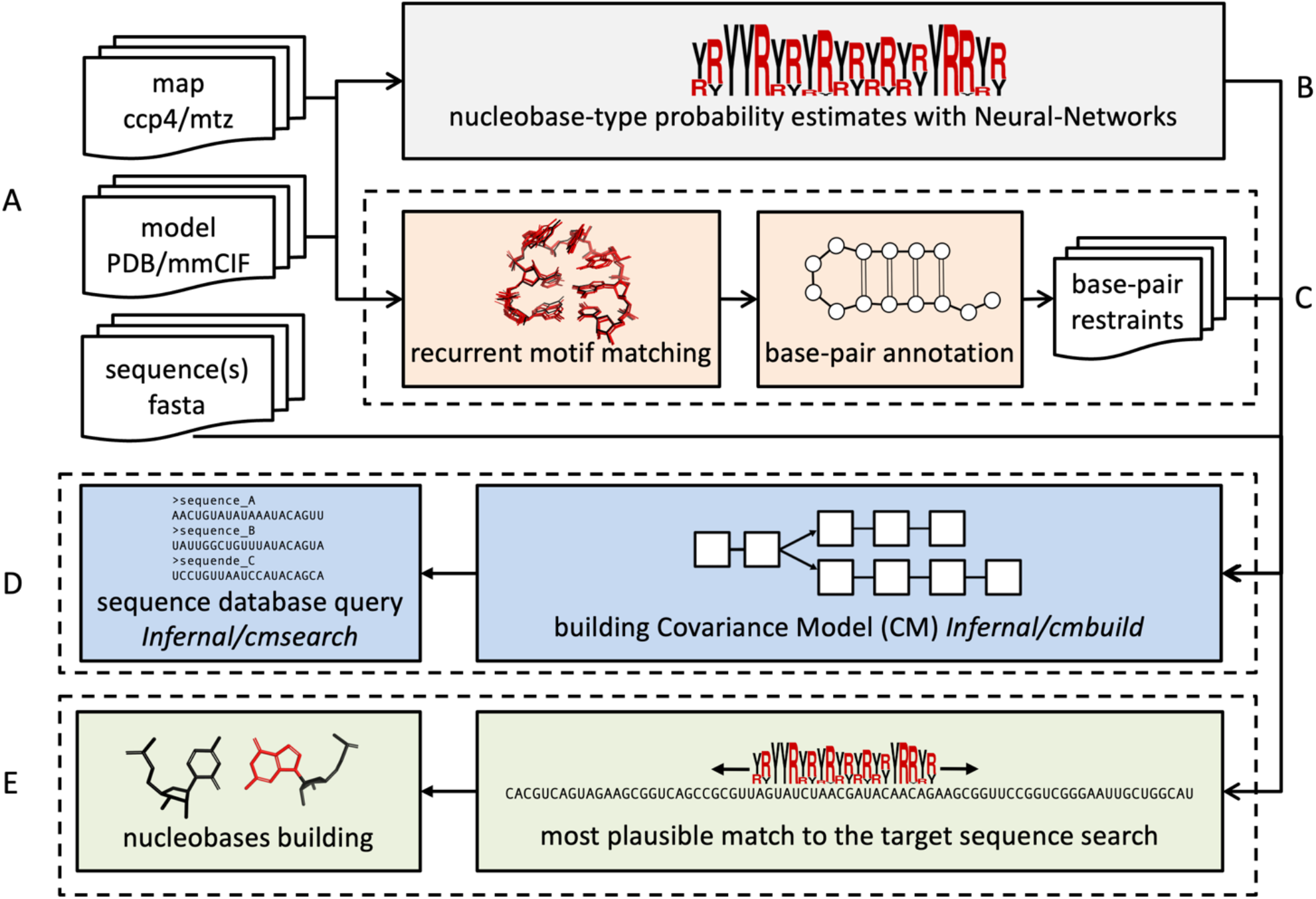
Schematic representation of the *doubleHelix* workflow. Key steps are color-coded and grouped in dashed rectangles; (A) input map (cryo-EM or MX), nucleic acid model, and target sequences (B) nucleobase-type probability estimation, (C) base-pair and refinement restraints assignment based on matched recurrent structural motifs, (D) sequence identification and (E) assignment based on estimated nucleobase-type probabilities and secondary structure. All steps are integrated in the software and performed automatically.

### Identification of unknown sequences of nucleic-acid models

For a nucleic acid model and a corresponding map (CCP4/MRC or MTZ formats for cryo-EM and MX respectively) program identifies the most plausible sequence from a sequence-database in FASTA format given estimated nucleobase-type probabilities and input model secondary structure (Figure 1D).

### Assignment of nucleic acid models to known target sequences

For a nucleic acid model and a corresponding map (CCP4/MRC or MTZ formats for cryo-EM and MX respectively) program assigns continuous polynucleotide chain fragments to the target sequence and rebuilds the bases accordingly. Apart from the estimated nucleobase-type probabilities, base-pairs identified within the fragments are used as additional restraints (Figure 1E).

### Sequence assignment validation in nucleic-acid models

Given a nucleic-acid model, a corresponding map (CCP4/MRC or MTZ formats for EM and MX respectively) and the set of all model sequences, the program evaluates the plausibility of the model’s sequence assignment. This feature is implemented as an extension of the *checkMySequence* program and uses an algorithm described previously (19).

## MATERIAL AND METHODS

### Recurrent structural motifs in nucleic acids

The *doubleHelix* program identifies base pairs in RNA and DNA models from a local similarity of backbone coordinates with small “search-fragments” of known secondary structure. Model, double helical A-RNA and B-DNA search-fragments were generated using the X3DNA suite (32). Non-helical search fragments were selected using RNA Bricks (33) database. Selected sets of recurrent RNA fragments classified as “loops” with at least 500, 100, 25, and 10 occurrences in the database correspond to 83, 532, 1430, and 2664 search-fragments respectively (as of 28 March 2020).

### Ribosome crystal structures for secondary structure assignment benchmarks

As a reference for the secondary structure assignment benchmarks, I used crystal structures of ribosomes available in PDB as of 28 March 2020. From all structures determined at a resolution better than 3.0Å I selected ones with crystallographic R-free factor below 0.3. To reduce the set redundancy, from each group of similar structures (e.g. originating from the same publication) I selected models with the lowest R-work/R-free difference. The resulting set contained eight structures originating from *Haloarcula marismortui, Thermus thermophilus*, and *Deinococcus radiodurans* (PDB entries 1s72, 4ybb, 7rqa, 1hnx, 1fjg, 1k73, 4y4o, 6oxi). For each of the models, secondary structure was determined using the ClaRNA program (34) and used as a ground-truth.

### Reference set of ribosome cryo-EM structures for sequence identification benchmarks

From the PDB I selected cryo-EM structures of ribosomes determined at a resolution better than 3.5 Å. Among 102 such structures available as of 4 February 2020 I selected 17 with half-maps available for download in the Electron Microscopy Data Bank (EMDB). For each of the half-map pairs local resolution maps were calculated using Resmap version 1.1.4 (35) with default parameters.

The selected models (PDB entries 3j79, 3j7a, 3j7q, 5iqr, 5mdv, 5mdw, 5mdy, 5ngm, 5umd, 5wdt, 5we4, 5wfs, 6okk, 6p5i, 6p5j, 6p5k, 6p5n) originated from five different organisms: *Plasmodium falciparum, Escherichia coli, Staphylococcus aureus, Sus scrofa* and *Oryctolagus cuniculus*. For each of them nucleotide sequences corresponding to RNA features annotated based on the genome sequence were downloaded from NCBI (36) and used as references for the sequence identification benchmarks. The reference sets contain tRNA, rRNA and ncRNA sequences, except for those corresponding to eukaryotic organisms that additionally contain mRNA sequences. To ensure that exact matches are available in the reference sets I added target rRNA sequences to each of them.

### Structures used for training neural network classifiers

As map features observed in cryo-EM and MX maps differ in fine detail, two separate neural networks were trained for each of these experimental methods.

For training the cryo-EM nucleobase-type classifier from the cryo-EM structures of ribosomes initially selected for sequence identification benchmarks, for which half-maps were not available in EMDB, I randomly selected 10 (PDB entries 5afi, 5mmi, 5u9f, 5wdt, 6eri, 6h4n, 6ogi, 6om0, 6q8y, 6sgc). Additionally, 142 PDB-deposited cryo-EM structures containing a DNA, but not an RNA component determined at resolution 3.5 Å or better with map-to-model correlation coefficient above 0.8 as estimated for complete models (including non-NA components) using phenix.map_model_cc (37) were added to the training set.

For training a crystal structure nucleobase-type classifier I selected eight structures and corresponding maps that were also used for benchmarking secondary structure assignment procedure. Moreover, 100 crystal structures randomly selected from PDB that contain a DNA, but not an RNA component, determined at resolution 3.5 Å or better with R-free - R-work below 0.3 were added to the training set.

### Ribosome crystal structures for sequence identification and assignment benchmarks

For RNA sequence identification benchmarks, I arbitrarily selected two crystal structure models of *Thermus thermophilus* 30S ribosomal subunit determined at a resolution 2.8Å (PDB entry 2uub) and 3.3Å (PDB entry 6mpi). For both targets re-refined structures were downloaded from the PDB_REDO server (38). Initially, a randomly selected 90% of ribosomal RNA nucleobases were mutated in both models (purines to pyrimidines and vice versa) keeping canonical base-pairing interactions identified using doubleHelix (the procedure is implemented in *doubleHelix* program and can be enabled with an option “*--randomize=0*.*9*”). The model coordinates were subsequently randomised with 0.2Å RMSD ignoring any geometry restraints and automatically refined using the PDB_REDO web server. For both models the randomisation procedure clearly affected the R-work/R-free factor values that increased from 0.19/0.23 to 0.25/0.29 and from 0.20/0.25 to 0.23/0.27 for better and worse resolution structures respectively. The automatically refined randomised models with corresponding maximum likelihood, combined 2mFo-DFc maps were used for sequence identification and assignment benchmarks.

For the sequence identification benchmarks nucleotide sequences corresponding to RNA features annotated based on the *Thermus thermophilus* genome were downloaded from NCBI and used for making queries.

### Neural network base-type classifier

To estimate the probability that a given nucleotide fitted into a map corresponds to a purine or pyrimidine two independent neural-network classifiers were prepared. The classifiers have identical architecture but are trained on distinct training sets derived from crystal structures or cryo-EM models and their respective maps.

Nucleotides are described with a vector of map values sampled around a putative base moiety (a residue descriptor). The map is sampled on a regular grid with 1.0 Å spacing. The grid is centred at the N1 or N9 atom for purine and pyrimidine respectively and spanned by orthonormal vectors defined by glycosidic bond (e_x_), the normal vector of the ribose best-fitting plane (e_y_), and their cross product (e_z_=e_x_ x e_y_). For a given nucleotide the input to the classifier contains a cloud of 403 grid points that are within 1.0 Å distance from any atom of a nucleobase mutated to Guanine in any rotation around the glycosidic bond. In practice a precomputed cloud is aligned to each nucleotide using C2’, C1’, and O4’ ribose or deoxyribose atoms.

The neural-network model input is a vector of length 403 (the residue descriptor described above). The model contains two, fully connected hidden layers. The first layer has a ReLU (Rectified Linear Unit) activation function, which sets all negative neuron inputs to zero, and 403 output features. The second layer has 2 output features and uses the log-softmax normalisation function enabling estimation of output classification probabilities. To avoid overfitting, an additional dropout layer was inserted between the two hidden layers. The dropout layer at each training step disables neuron connections selected at random with probability *p*. The models were trained for 1,000 epochs with *p*=0.5, a batch size of 20 residue descriptors in each parameters update cycle, and a 10% validation set. The models were trained using the ADAM optimization algorithm (39) with a learning rate of 1e-5 that resulted in the best test-set accuracies.

For training the crystal structure classifier I used 84,887 and 9,431 nucleobase descriptors for training and test-set respectively. The accuracies of a resulting model were 0.98 and 0.96 for the training and test sets, respectively. Similarly, for training the EM classifier I used 85,092 and 9,454 residue descriptors in training and test-sets, respectively. The resulting model estimated accuracies were 0.95 and 0.92 for training and test set, respectively.

### Secondary structure assignment procedure

Unlike DNA, which occurs in nature predominantly in a double-helical form, RNA molecules are often single-stranded and fold into complex structures stabilised by stacking and base-pairing interactions (40). Folded RNA molecules have a modular architecture in which the double-helical regions are intertwined with different types of loops that define the topology of the structure and stabilise it through long-range interactions. Many of these loops are recurrent and can be found in a similar structural context in many, possibly evolutionary unrelated, RNA molecules (33). Most importantly, it is the overall module geometry and base-pairing pattern rather than the nucleotide sequence that is conserved across different occurrences of the same module (41). This feature of RNA molecules is used in the doubleHelix program for the inference of base-pairing interactions from the local geometry of sugar-phosphate backbone. This approach ignores both identities and mutual orientation of bases, which is particularly important in the analysis of preliminary, not fully refined models, where base identities are not yet known and their coordinates, unlike relatively heavy backbone, may be inaccurate.

The program superposes small RNA or DNA search-fragments of known secondary structure onto the input model using an algorithm described previously as a part of a model-building program *Brickworx* (2). First, all possible triplets of phosphate group P-atoms in a search-fragment are structurally aligned with similar P-atom triplets from the input structure. Resulting rigid body transformation is applied to the complete fragment to identify matching nucleotides in the search fragment and input structure. Finally, the match is refined using all sugar-phosphate backbone atoms. If the resulting root-mean-square deviation (RMSD) is below 1.0Å (the threshold defined in the Results section), search-fragment base pairs are assigned to the corresponding residues in the input model. If multiple, overlapping matches of search-fragments are identified, the one with the lowest RMSD is selected.

For the sake of computational efficiency, the input model processing is divided into two steps. Firstly, only a double-helical fragment is matched to identify Watson-Crick base-pairs. In this step, A-RNA or B-DNA search fragments are used depending on the target nucleic acid type. Next, all nucleotides within stacked Watson-Crick base-paired regions (except flanking residues) are removed from the input model. In the second step, used only in case of RNA targets, a predefined set of recurrent RNA motifs is matched to the remaining nucleotides in the input model. In case of base-pair assignment conflicts, the ones detected in the second step are given preference.

### Sequence identification procedure

For the identification of the most plausible sequence in a database, given input model residue-type probabilities and secondary structure *doubleHelix* uses sequence-comparison tools from the INFERNAL suite (42) (Figure 1D). Initially, predicted residue-type probabilities are converted into a multiple sequence alignment (MSA), where fractions of residue types in each column correspond to predicted probabilities. The MSA, combined with base-pairing pattern is encoded in a Stockholm format (STO) which is an input to the *cmbuild* program. The resulting Covariance Model (CM) is further calibrated with *cmalibrate* and used to query sequence databases using the *cmsearch* program with default parameters. The Hidden Markov Model (profile-HMM) queries are enabled by adding the *-- hmmonly* keyword to *cmsearch* and skipping the calibration step. Sequence hits with the lowest E-values are returned to the user (3 by default).

### Sequence assignment procedure

Analogously to the sequence identification procedure, *doubleHelix* uses the estimated base-type probabilities to assign RNA or DNA models to a target sequence. For a continuous polynucleotide fragment in the input model a neural network classifier is used to estimate base-type (purine or pyrimidine) probabilities for each residue. The resulting scoring matrix is aligned to the target sequence and the probability of each tentative alignment is approximated by a product of the probability estimates for each residue in the fragment, assuming their independence. If any residue pair in the fragment forms a Watson-Crick base-pair (detected using a procedure described above) an additional, low-probability correction factor (0.1) is used if for a tentative alignment the two bases are either both purines or pyrimidines, which is very unlikely for a Watson-Crick interaction. Otherwise, the correction term is 1.

Although the neural-network classifier has been calibrated and the predicted residue-type probabilities generally reflect expected frequencies, the accuracy of predictions may vary depending on local map resolution and quality of the models (11). Therefore, for each tentative assignment of a fragment to a target sequence the method estimates a p-value, or probability that a given alignment has been observed by chance. A tentative alignment probability is compared to a background distribution of the fragment alignment probabilities for a long, random sequence. To additionally account for the varying target-sequence lengths an additional extreme-value distribution theorem correction is applied as described before (19).

### Implementation

The *doubleHelix* program was implemented using Python3 with an extensive use of numpy (43), scipy (44), CCTBX (45) and CCP4 (46) libraries and utility programs. The neural network classifier used in this work was trained using Pytorch (47) and deployed to a C code using keras2c (https://github.com/f0uriest/keras2c). For making the rRNA sequence database queries the program uses INFERNAL suite version 1.1.4.

## RESULTS AND DISCUSSION

### Secondary structure assignment

The base-pair assignment in the *doubleHelix* program relies on structural alignment of recurrent motifs with known secondary structure (search-fragments) to the input model. The method uses five different sets of search-fragments. First, A- or B-form double helices, for RNA and DNA target models respectively, are tried. Additionally, for RNA models, the method uses four sets of the most frequent recurrent RNA motifs from the RNA Bricks database (see Materials and Methods for details). Although the double-helical search fragments can be used for the identification of the canonical Watson-Crick base-pairs only, the other search fragments allow for the identification of any other base-pairing interaction type.

The base-pair assignment procedure parameters providing maximum performance are 2-base-pair double-helical search fragments with 1.0Å RMSD threshold (Figure 1A). This can be explained with a relatively high structural heterogeneity that was observed in double-helical structures (48) that cannot be represented using longer, idealised search models. Interestingly, an additional search step with recurrent RNA motifs with at least 500 occurrences in the RNA Bricks database further improves precision of the Watson-Crick base-pairs assignment. The resulting Watson-Crick base-pair assignment f1-score 0.94 is comparable to a value of 0.95 reported for a recently described program CSSR (49), which also relies on backbone conformations only. With *doubleHelix* this translates to the recall of 0.91 and precision 0.98 (no corresponding results were reported for CSSR). Unlike doubleHelix, however, CSSR focuses exclusively on pairs of nucleotides compatible with canonical base pairing (A/U, G/C, or G/U) that makes the two methods not directly comparable. Another advantage of the doubleHelix approach over CSSR is its ability to identify stackings and non-canonical base-pairs with recall/precision of 0.63/0.89 and 0.47/0.94 respectively (Figure 2B). This, however, requires the use of a larger set of recurrent RNA motifs with at least 100 instances in the RNA Bricks database that results in an increased computational cost. For example, for a tRNA model (76 nucleotides, PDB entry 1ehz) processing time on a standard laptop increases from 4 seconds in default configuration to 13 seconds. For a complete porcine 28S rRNA (3938 nucleotides, PDB entry 3j7q) these times increase from 1 to almost 8 hours. This, however, is needed only for the accurate assignment of non-canonical base-pairs in RNA models (e.g. as refinement restraints). By default, for the identification of Watson-Crick base-pairs the *doubleHelix* uses the smallest set of RNA recurrent motifs providing maximum performance at a reasonable computational cost.

**Figure 2.**
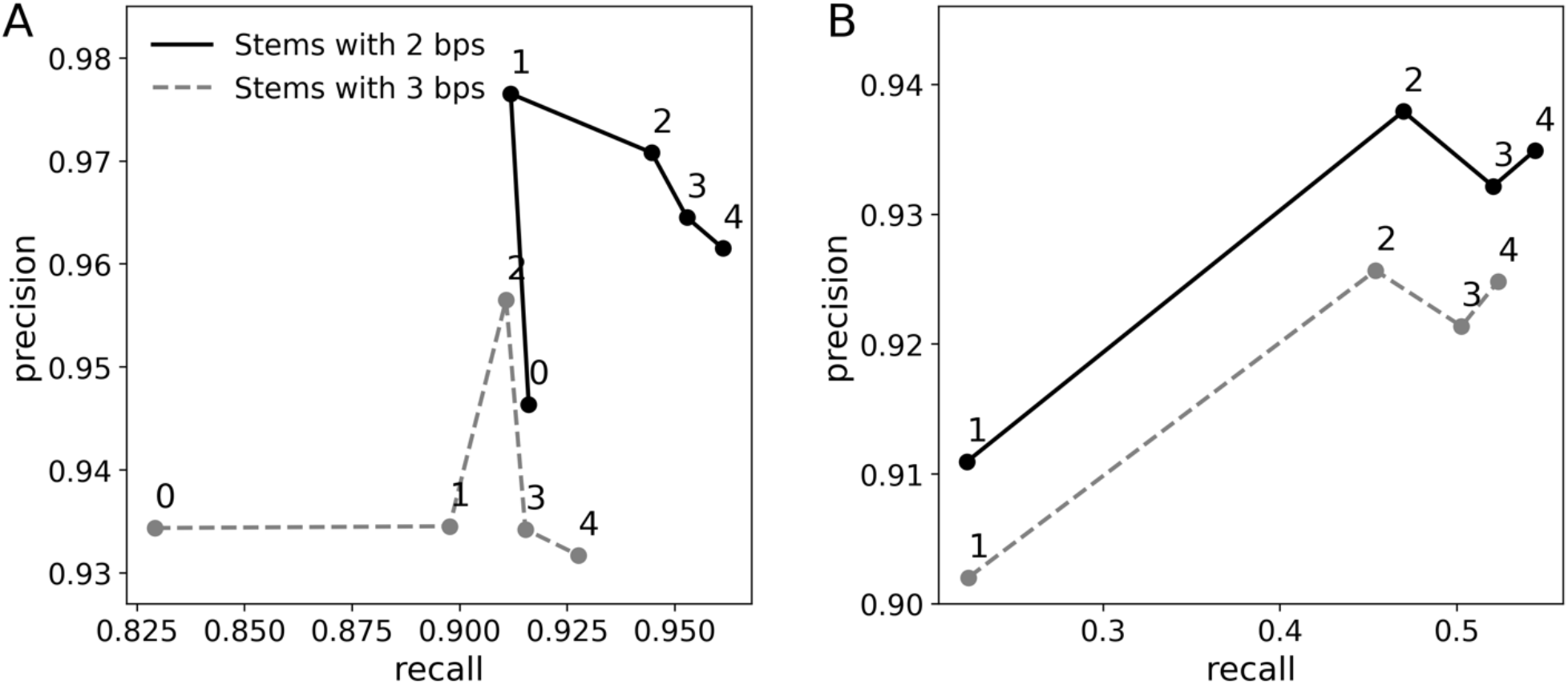
Performance of RNA secondary structure assignment based on structural alignment of recurrent motifs for (A) canonical (Watson-Crick) and (B) non-canonical base-pairs. Each data point represents performance for a given stem length (2 or 3 base pairs) and main-chain atoms RMSD maximising classification f1-score (harmonic mean of precision and recall). Data-point labels correspond to the number of recurrent RNA motifs used for model interpretation; RNA stem only (0), motifs with at least 500 (1), 100 (2), 25 (3) and 10 (4) occurrences in RNA Bricks database. Precision and recall were estimated using ClaRNA (34) base-pair classification as ground truth.

### Sequence identification in cryo-EM

For the identification of a nucleic acid model the doubleHelix program finds the most plausible sequence in a database given nucleobase-type probabilities (purine or pyrimidine) estimated based on a backbone model and corresponding cryo-EM map. Secondary structure restraints, derived directly from a backbone model, are used as an additional source of information. Both base-type probabilities and base-pairing information are used to query sequence databases using the INFERNAL suite as described in the Materials and Methods section. I observed that this approach allows for a sequence identification up to 4.5Å local resolution for fragments of 50 amino acid residues (Figure 3A) when Covariance Models (CMs) and secondary structure information is used. The use of Hidden Markov Models (HMMs), which neglects base-pairing information, clearly reduces the method performance (Figure 3A). The use of longer fragments of 100 residues, further increases the resolution limit of the method applicability up to 5.5Å when the base-pairing information and CMs are used (Figure 3A). Overall, the E-value provided by the INFERNAL suite is a reliable measure of the sequence identification reliability. There are, however, a few model fragments for which the identified sequence has a relatively low sequence identity to the target, despite a reliable E-value score (Figure 3B). This issue can be attributed to the reduction of the sequence identification problem to purines and pyrimidines only that may occasionally make different sequence fragments practically indistinguishable.

**Figure 3.**
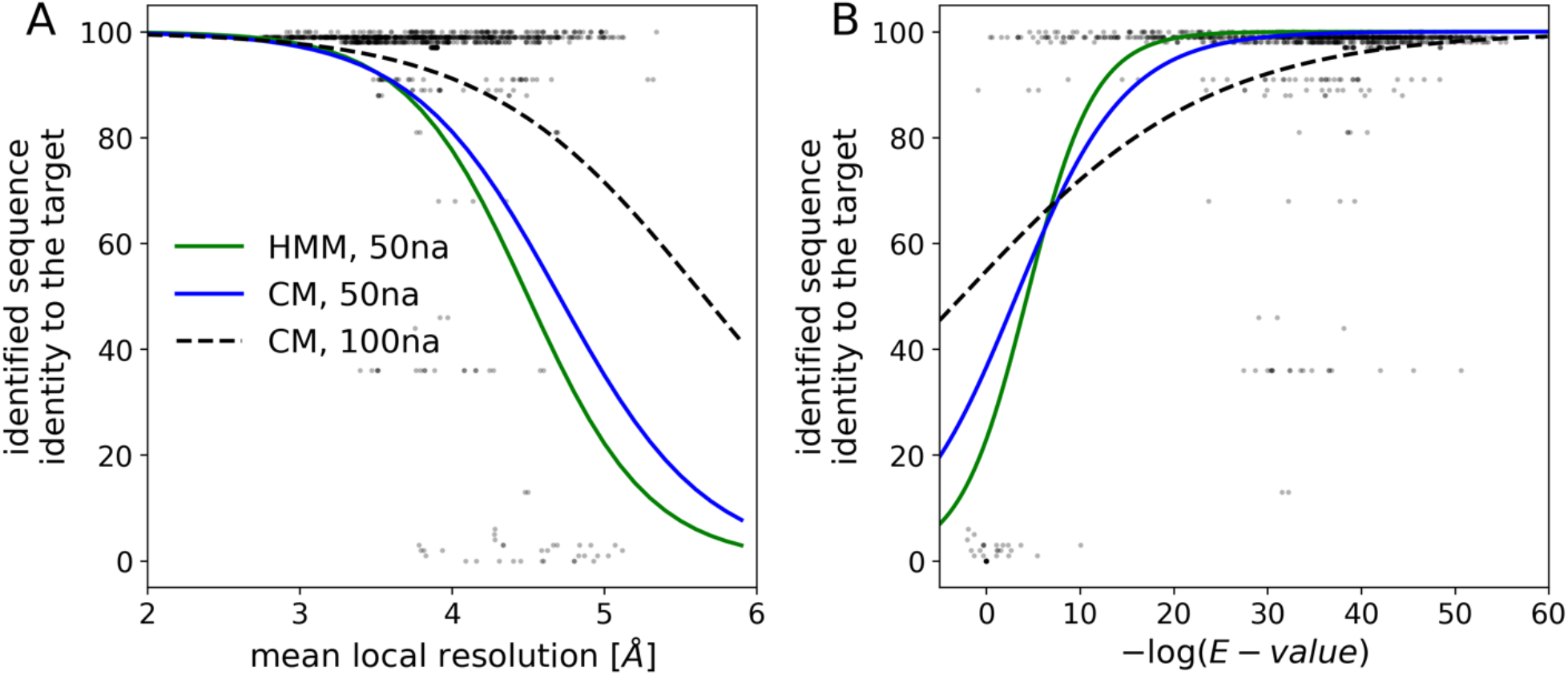
Sequence-identification benchmarks with continuous fragments of 50 and 100 nucleic acid residues selected at random from cryo-EM ribosomal RNA models. Comparison of the method performances for an identification of model sequence with (CM) and without (HMM) the use of base-pairing information. The sequence identification performance is shown as a function of (A) local resolution of EM maps and (B) E-value of the sequence assignment estimated by the INFERNAL suite. The (B) plot horizontal axis shows -log(E-value); higher values correspond to lower E-values and more reliable sequence identification results. The continuous curves are logistic regression estimates of a probability that an identified sequence will have at least 90% sequence identity to the target sequence.

### Sequence assignment in cryo-EM

It has been shown that a neural network-based assignment of protein model sequence can provide reliable results up to local map resolutions where *de novo* model building would be very challenging (50). Moreover, a carefully derived sequence assignment reliability score, the p-value, accurately separates correct from wrong alignments independently of local map resolution. The assignment of polynucleotide sequences, however, presents a particular challenge compared to proteins. First of all, nucleic acid models are often built *de novo* into lower resolution map regions, as double-helical fragments are excellent and universal models for map interpretation. Moreover, even at moderate resolutions the target sequence is effectively reduced to only two types of nucleobases (purines and pyrimidines) that greatly increases sequence assignment ambiguity.

I observed that similarly to proteins the neural-network classifier implemented in doubleHelix provides a reliable means of assigning polynucleotide sequences to model fragments at local resolutions as low as 5Å, even though the required fragment lengths are clearly longer (Figure 4A). For a total number of 18,655 RNA test-fragments of 20 residues (see Materials and Methods for details), the assigned sequence matched the corresponding model in 83% (15,461) of cases. For longer fragments of 40 residues the number of correctly assigned sequences increases to 95% (17,383).

**Figure 4.**
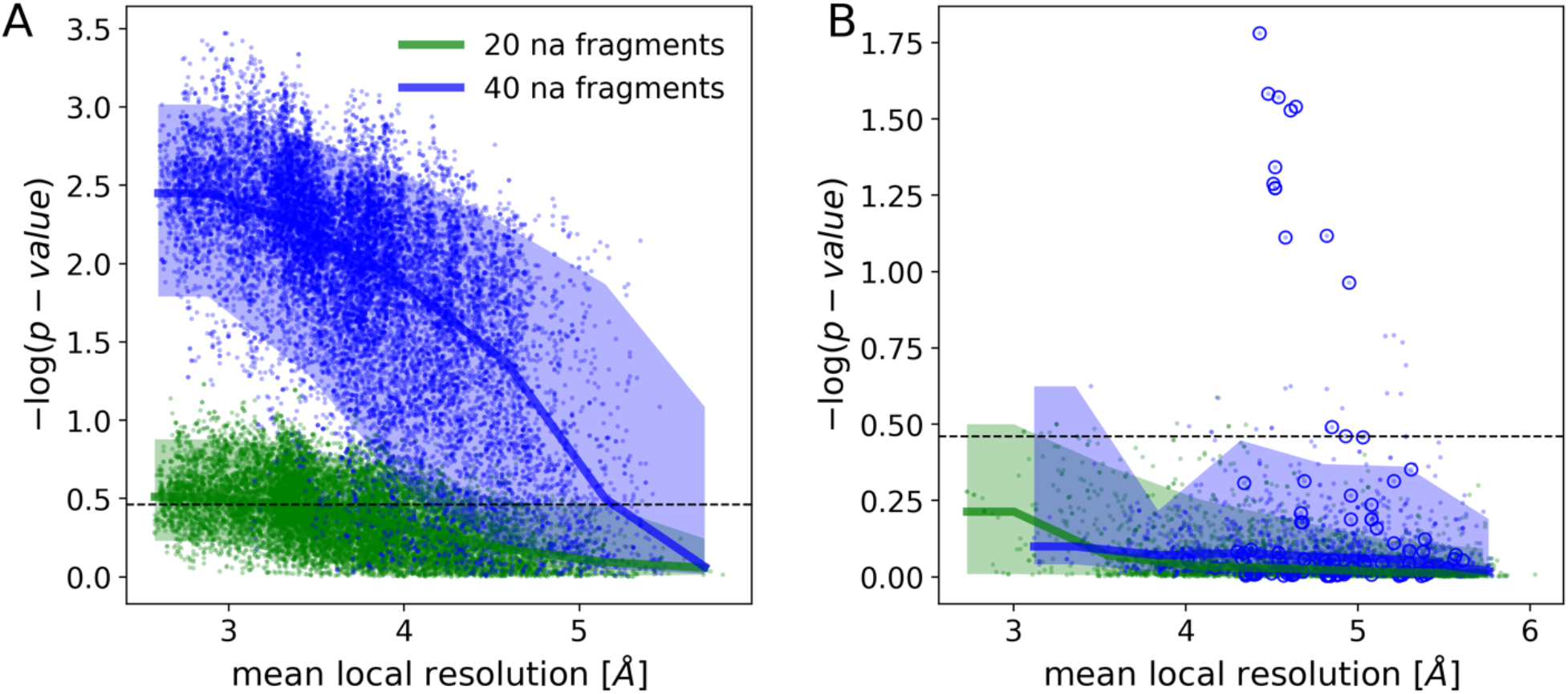
Medians and 90% confidence intervals for sequence assignment p-value as a function of local resolution for RNA chain test-fragments of 20 and 40 nucleic acid residues. Panel (A) shows fragments with sequences matching reference. Fragments for which assigned and reference model sequences differ are presented in panel (B). The dashed line corresponds to a 99.5% one-sided confidence interval estimated for fragments of 20 nucleic acids with an assigned sequence different from the input-model sequence. Blue circles depict an outlier presented in the text (porcine 28S rRNA, PDB entry 3j7q). The plots’ ordinate axes show –log(p-value) for the test-fragments; higher values correspond to lower p-values and more reliable sequence assignments.

The p-value, or the probability of observing a given sequence assignment by chance, provides a reliable estimate of the alignment accuracy that doesn’t depend on local map resolution (Figure 4B). Indeed, a one-sided 99.5% confidence interval for fragments with sequence assignment that doesn’t match the reference (dashed line on Figure 4) corresponds to 62% and 17% of cases with matching sequences for fragments of 20 and 40 residues respectively. Although sequence mismatches with a p-value outside this region are expected to be very rare, I observed several such test-fragments in the benchmark set (blue circles on Figure 4B), all of them correspond to a single model of porcine 28S rRNA (PDB entry 3j7q). I will discuss this outlier in more detail in the next section.

### Sequence assignment outlier: mammalian 28S rRNA

In the cryo-EM benchmark set, I identified several clear outliers, where RNA test-fragments were assigned reliable (low p-value) sequences different from the reference model (Figure 4B). All the fragments originate from an expansion segment (ES7a) of a cryo-EM structure of porcine 28S ribosomal subunit determined at 3.4Å resolution (PDB entry 3j7q). Closer inspection of the model revealed several nucleobases poorly fitting the EM reconstruction, which is understandable given limited resolution, but no clearly visible sequence assignment issues. As there is no higher resolution structure for the porcine ribosome available, which could be used to validate the sequence register, I decided to use as a reference the closest homologue from rabbit (98% of sequence identity), for which a structure determined at 3Å resolution has been recently deposited (PDB entry 6r5q). For a detailed comparison I selected a fragment of the ES7a that has a strictly conserved sequence in both organisms according to an alignment generated using R-coffee (51). Structural alignment of the corresponding model fragments, however, revealed multiple differences between corresponding nucleobase identities, several of them resulting in base-pairing violations (Figure 5C). I also observed that several differences in sequence preserving secondary structure are visible as clear density-fit outliers (Figure 5A). The problem is easily solved by shifting the sequences of the two chain fragments by one residue, as suggested by the *doubleHelix* program, which results in a perfect fit between the porcine and rabbit models (Figure 5B).

**Figure 5.**
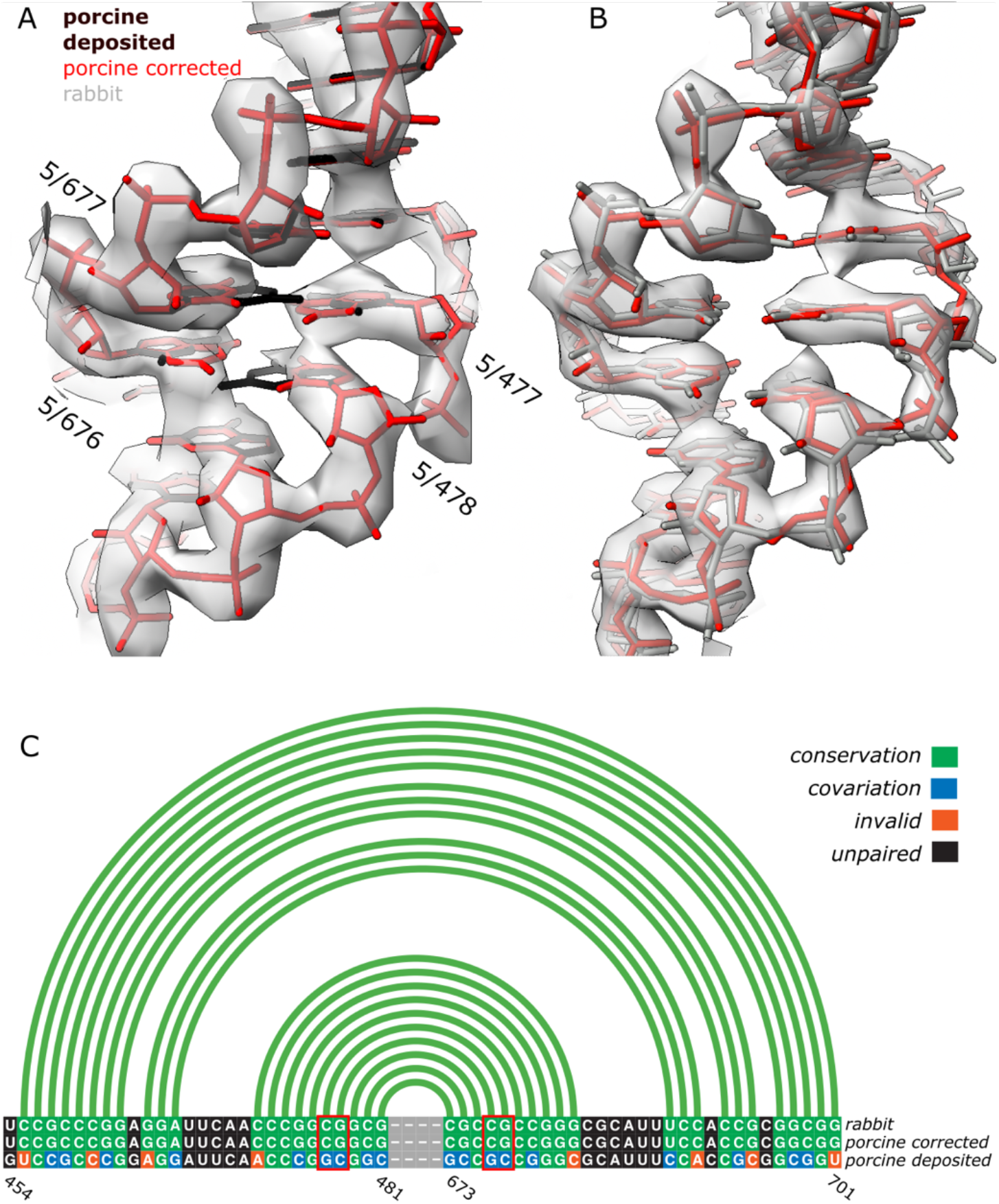
Fragments with strictly conserved sequence of porcine (A) and rabbit (B) expansion segments (ES7a) in 28S rRNA with corresponding cryo-EM reconstructions at 3.4Å and 3.0Å resolution, respectively. Black model on panel (A) and grey on panel (B) represent deposited coordinates whereas porcine structure with sequence re-assigned using *doubleHelix* is depicted in red. Aligned sequences and secondary structures of the rabbit and porcine models are presented on panel (C). Although the register shift in deposited porcine structure preserves most of the base-pairs (green and blue boxes), several of them are visible as clear density-fit outliers (bases in red rectangles and labelled on panel A). There are also multiple secondary structure violations (shown in orange). Secondary structure presented on panel (C) was identified from the corrected model using ClaRNA. The figure was prepared using ChimeraX (52) and R-chie webserver (53).

### RNA sequence identification and assignment in MX

A crystallographic diffraction experiment provides only amplitudes of complex structure factors required for calculating electron-density maps. Missing phases need to be derived from other sources, for example a tentative model of the unknown crystal structure from Molecular Replacement procedure. The use of model-derived phases for calculating electron-density maps inevitably results in so-called “model bias” - the presence in a map of model features that are absent in a crystal structure. The same issue may be expected when a tentative crystal structure model polynucleotide sequence doesn’t match an unknown crystal structure. Although the model bias is reduced in maximum likelihood maps (54), routinely used for model building and refinement in MX, the problem is not completely eliminated. To investigate how the model-bias issue affects the sequence identification and assignment procedures I benchmarked them using ribosome crystal structures with randomised rRNA sequences. I used two crystal structure models of *Thermus thermophilus* 30S ribosomal subunit determined at resolutions 2.8Å (PDB entry 2uub) and 3.3Å (PDB entry 6mpi). Additionally, to remove any effect related to the presence of protein chain models that were refined in the presence of the correct rRNA sequences all atomic coordinates were randomised with 0.2Å RMSD. The resulting, randomized 30S models were refined using REFMAC5 and the PDB_REDO webserver. Interestingly, observed model bias in both randomised structures is moderate and clear sequence mismatches can be noticed for few (but not all) nucleobases (Figure 6B and 6D).

**Figure 6.**
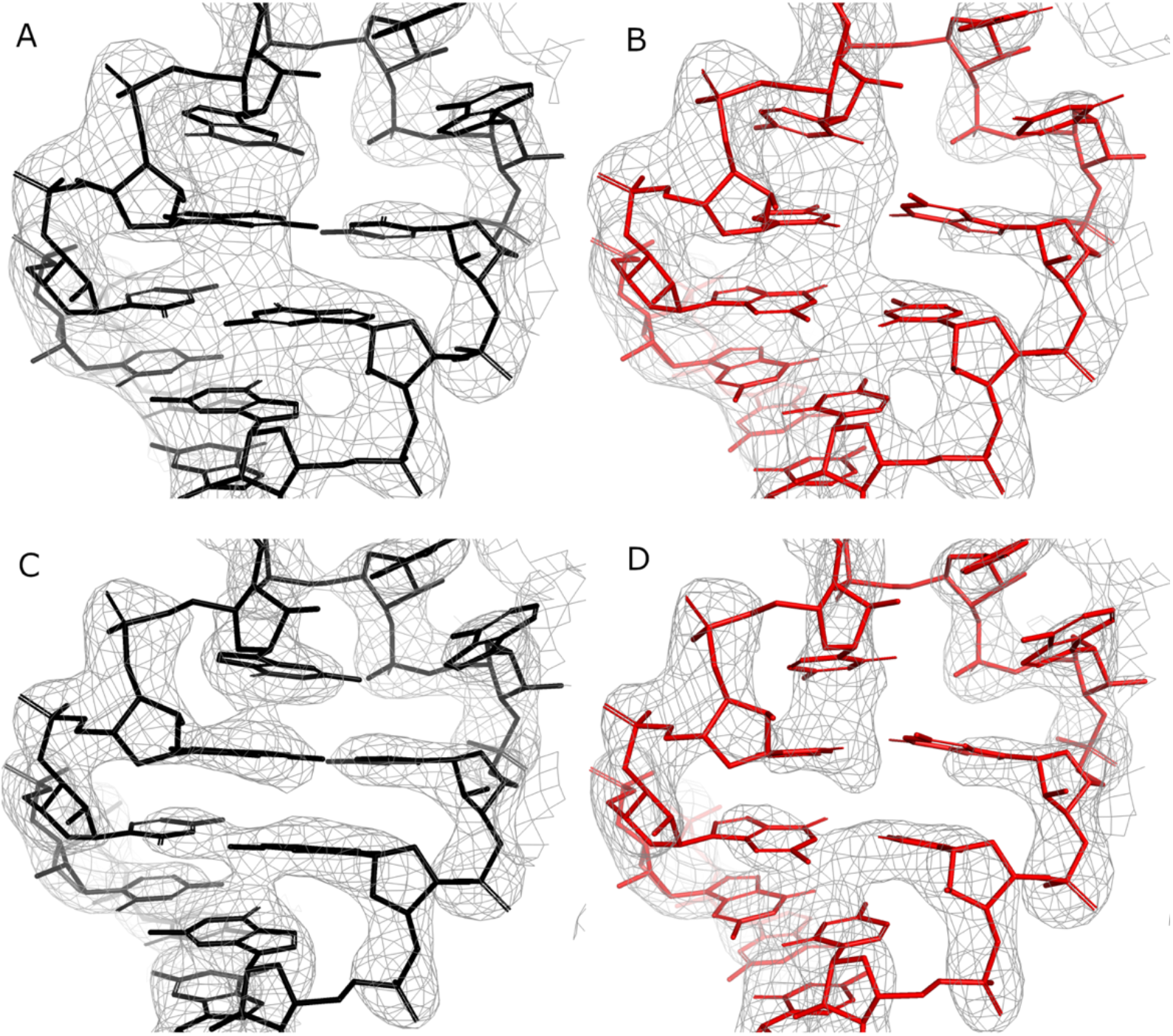
Corresponding fragments of *Thermus thermophilus* 28S ribosomal subunit crystal structures used for the sequence identification and assignment benchmarks. Two models with randomised rRNA sequence were generated based on crystal structures determined at a resolution of 2.8Å (PDB entry 2uub, panels A and B) and 3.3Å (PDB entry 6mpi, panels C and D). The panels depict residue range 1262-1273 with corresponding maximum likelihood 2mFo-DFc maps contoured at 2***σ*** level; as deposited (A, C) and after randomising atomic coordinates and mutating 90% of nucleobases (B, D, see Materials and Methods for details). Both deposited and randomised structures were automatically refined using the PDB_REDO web server.

The sequence identification procedure was very effective for the randomised 2.8Å resolution model. Among 1,000 continuous, randomly selected rRNA fragments of 100 residues, doubleHelix identified a correct target sequence in 99% of cases. For shorter fragments of 50 residues this fraction reduces to 76%. At lower resolution (randomised 6mpi model at 3.3Å resolution) the performance clearly reduces. The program provided a correct hit in 86% and 62% and of cases for test-fragments of 100 and 50 residues respectively. Interestingly, I also observed that the use of secondary structure information for the sequence identification clearly improves the method performance. Without base-pairing restraints the fraction of correctly identified sequences for fragments of 100 residues reduces to 90% and 72% for better and worse resolution structures. In all cases the incorrect hits can be easily filtered based on E-value returned by INFERNAL suite where values smaller than 0.1 guarantee a correct solution (Figure 7A).

**Figure 7.**
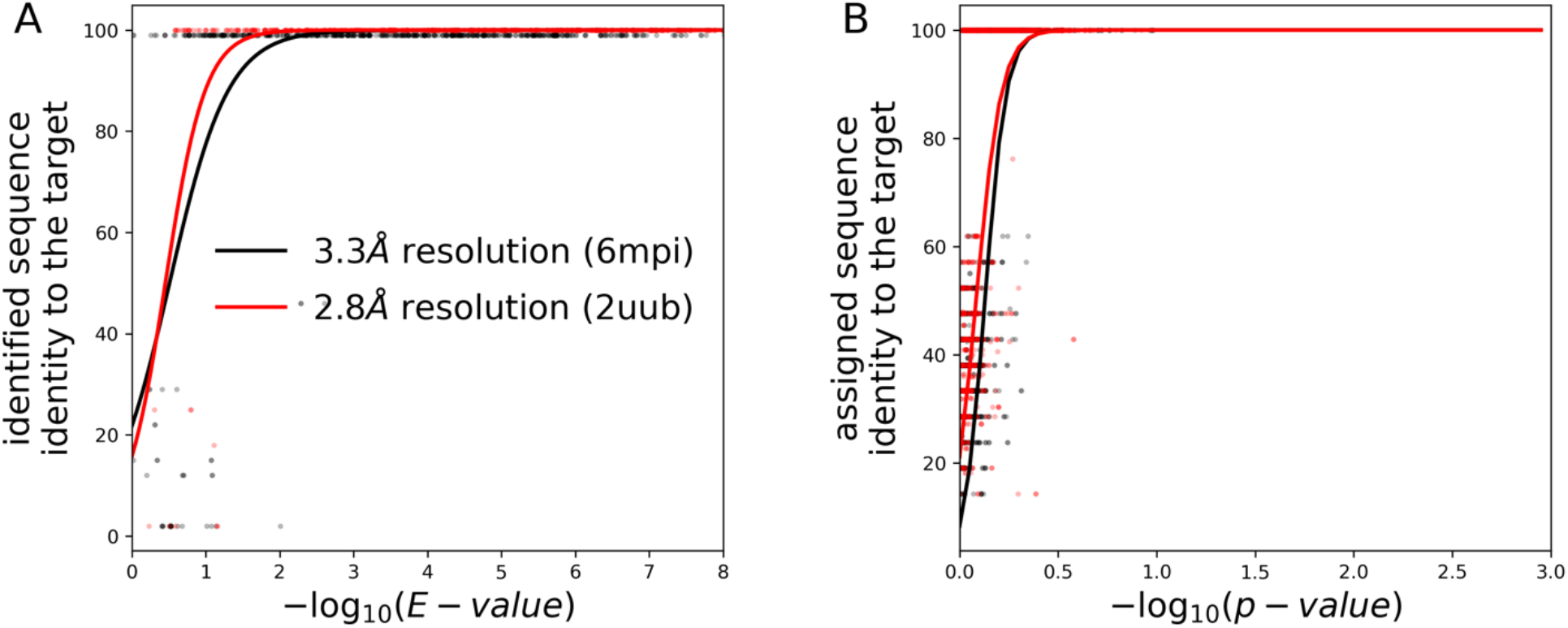
Performance of (A) sequence-identification and (B) sequence-assignment for fragments selected at random from two MX ribosomal RNA models determined at a resolution of 2.8Å (PDB entry 2uub) and 3.3Å (PDB entry 6mpi). The random fragments used for sequence identification and assignment had 50 and 20 nucleic acids, respectively. The continuous curves are logistic regression estimates of a probability that an identified sequence will have at least 90% sequence identity to the target sequence.

For sequence assignment the method correctly identified sequences of 46% and 65% of 1,000 continuous rRNA fragments of 20 nucleic acid residues selected at random from randomised models at worse and better resolution respectively. These numbers increase to 82% and 94% for longer fragments of 40 residues. The correct sequence assignments are also clearly separated from incorrect ones by p-value with the 99.5% one-sided confidence interval for wrongly assigned fragments matching a value estimated for EM models (Figures 4B and 7B).

### Case study: sequence register errors in crystal structure of the large ribosomal subunit of *S. aureus*

The integration of *doubleHelix* with previously developed *checkMySequence* program allows for a fully automated sequence assignment validation of complete protein-nucleic acid complexes. Benchmarks of the method revealed a particularly interesting structure of the large ribosomal subunit from *S. aureus*, where *checkMySequence* revealed a plausible register shift errors in two ribosomal proteins (L18 and L4) and 5S rRNA. The structure was determined at 3.5Å resolution (PDB entry 4wce,(55)) and refined to R-work/R-free factor values of 0.202/0.246 and *clashscore* 11.

A detailed discussion of the register shift error correction in ribosomal proteins is out of scope of this work and will be presented only briefly. The protein chains were replaced with corresponding predictions from AlphaFold2 database (release 4 for UniProt entries Q2FW07 and Q2FW22 for L4 and L18 respectively) subsequently refined using COOT in real space with self-restrains generated at 5Å cut-off. Both predicted models were assigned a very high confidence score (pLDDT>90), which was observed to usually correspond to minor differences in loop and side-chain conformations compared to reliable experimental models (56). Comparison of the deposited and corrected protein chains of both proteins revealed multiple plausible tracing issues and confirmed register-shifts suggested by *checkMySequence*.

The target sequence of the 5S rRNA chain consists of 114 residues that were all traced in the map (chain Y in the deposited model). The *checkMySequence* scan revealed an alternative sequence assignment with p-value of 0.01 (very reliable according to Figure 7B) for a chain fragment following residue 81. The alternative sequence corresponds to a register shift by one base which suggests that a residue could have been omitted in the deposited model (deletion). Although closer inspection of the crystal structure revealed several clear density outliers for bases (Figure 8A), there were no signs of tracing issues. To confirm the chain sequence correctness, I performed sequence identification using *doubleHelix* with the deposited 5S chain coordinates, corresponding maximum likelihood 2mFo-DFc map, and a set of RNA sequences for *S. aureus* strain NCTC8325 downloaded from NCBI. At the time of writing (25.11.2022) there were two assemblies available (GCF_900475245 and GCF_900475245) each containing roughly 100 tRNA, rRNA, and ncRNA sequences.

**Figure 8.**
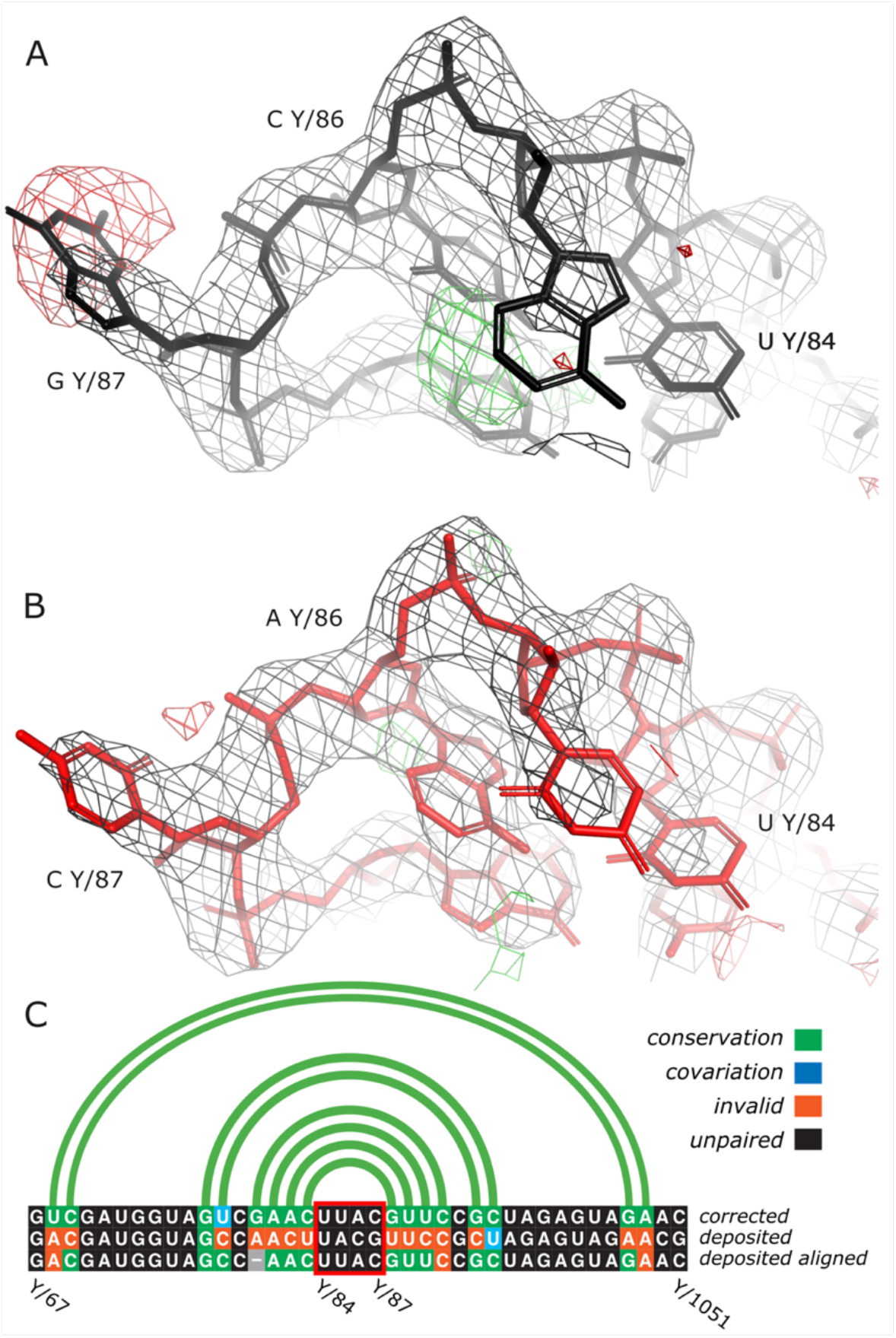
Fragment of 5S rRNA model from a crystal structure of large ribosomal subunit of *S. aureus* refined at 3.5Å resolution. High negative (red) and positive (green) difference density map values near Guanine Y/87 and Cytosine Y/86 correspond to excess and missing atoms in the model (A). After re-assigning the model to a sequence of different 5S gene variant and subsequent restrained refinement in REFMAC5 and PDB_REDO map-model fit clearly improves (B). The new sequence also clearly improves base-pairing pattern of the model (C). For clarity, only a sequence fragment including the stem loop depicted in panels (A, B) is shown (highlighted with a red box). Most sequence mismatches between the two 5S sequence variants can be corrected by introducing a gap to the alignment (shown as a grey box) that corresponds to a single base register-shift identified by *checkMySequence*. Maximum likelihood combined 2mFo-DFc and difference mFo-DFc maps on panels (A, B) are contoured at 2***σ*** and 3***σ*** level respectively. Secondary structure presented on panel (C) was identified in the corrected model using ClaRNA. The figure was prepared using Pymol (31) and R-chie webserver (53).

In the RNA sequence sets *doubleHelix* identified two, different 5S genes (96% sequence identity), both very reliable hits with E-value below 1e-20 (see Figure 7A). One of the sequences, however, scored visibly better that the other (7e-25 versus 4e-21 for NC_007795.1_rrna_7 and NC_007795.1_rrna_6 respectively), which has been previously shown to be a good evidence for a better fit to the data for protein models (28). The deposited 5S model was assigned the sequence variant with worse E-value, which resulted in the map-model fit outliers mentioned above (Figure 8A). The model, after re-assigning the 5S sequence variant identified with better E-value and subsequent restrained refinement with REFMAC5 and PDEB_REDO shows much better fit to the map (Figure 8B). The new base identities are also more favourable for the formation of canonical base-pairs in the model (Figure 8C). The final model of complete ribosome, after correcting tracing issues in the two protein chains (L4 and L18) and the 5S chain sequence refines with clearly better scores; R-work/R-free reduces to 0.195/0.237 (from 0.202/0.247) and clashscore to 5 (from 11). The additional base-pairs, visually better map-to-model fit and reduced global model-quality scores together provide good evidence of improved agreement of the corrected model with the experimental data.

## CONCLUSIONS

Sequence assignment is a key step of macromolecular model building that may lead to difficult to identify errors affecting structure interpretation. Nevertheless, it has been shown that protein models deposited in the PDB, despite expensive model validation efforts, contain many register-shift errors (16,20,57). Validation and assignment of nucleic acid sequences presents a particular challenge compared to proteins; the models are usually poorly resolved in cryo-EM and MX maps and available sequence-information is in practice limited to two nucleobase-types. Moreover, validation of nucleic acid models addresses predominantly backbone conformations that are rarely affected by nucleobase-sequence assignment issues. In consequence, the reliability of available, experimentally determined nucleic acid models is very difficult to assess. This may result in unintended error propagation, where a newly deposited model contains an error inherited from an earlier one used for model building. Errors can also detrimentally affect efforts of bioinformaticians working on large-scale structural analyses or structure prediction methods.

Here I presented *doubleHelix*; a new program for a comprehensive assignment, identification, and validation of nucleic acid sequences in cryo-EM and MX. I show that the approach, which relies on a neural-network base-type classifier, can successfully identify and assign sequences of cryo-EM model fragments at local resolution as low as 5Å. I also show that base-pairing information derived from backbone-geometry clearly improves the program’s performance at lower resolutions but is not essential. This is particularly important for large nucleic acid structures with very low content of paired bases; for example the trypanosomal mitochondrial ribosome (58).

The *doubleHelix* software can be also used for generating base-pairing restraints ready-to-use with the most popular cryo-EM and MX model-building and refinement tools (REFMAC5, PHENIX, COOT). I have shown that the approach can successfully identify 91% of canonical base-pairs with 98% precision without relying on base conformation or identity. This may be particularly useful in early stages of the model building process as available base-pairing restraints generation approaches are strictly dependent on base-identities and the detection of hydrogen-bonding patterns that requires accurate relative positioning of paired-bases (LibG (59), Phenix suite (60)). An exception here is a pipeline implemented in PDB_REDO (61) relying on base-pair assignment by DSSR suite, which is by design less sensitive to the structure distortions (62).

The presented *doubleHelix* benchmarks revealed a plausible sequence-register error in an expansion segment ES7a of a mammalian ribosome model deposited in the PDB (PDB entry 3j7q/EMDB entry EMD-2650). Expansion segments (ES) are present only in eukaryotic ribosomes and exhibit a surprising level of variability between different organisms. Nevertheless, the function of ES remains poorly understood, which makes them an important research target (63). Ribosomes are usually highly conserved across all kingdoms of live and ribosome models already available in the PDB can be often used to greatly simplify model building and refinement process of newly determined structures. The high variability of the ES at a structural and sequence level makes them one of the few rRNA regions that require in-depth modelling. Not surprisingly, this results in sporadic errors, which may hinder efforts aimed at understanding ES function. The problem is even more important for newly determined nucleic-acid complex structures for which at least partial models that could be used for model building are not available in the PDB. This makes the *doubleHelix* program particularly useful in structure determination using cryo-EM and MX as a reliable tool for nucleic-acid model sequence assignment and validation.

To facilitate the use of *doubleHelix* for model validation, it has been incorporated into a previously released, open-source sequence-assignment validation tool *checkMySequence* that can now process complete protein-nucleic acid complexes. The method enables a conceptually simple and fully automated detection of the most common sequence assignment issues in models of proteins, RNA, DNA, and their complexes that include register-shifts, sequence mismatches (single-residue differences between model and target sequence), problems with residue numbering (e.g. continuous residue numbering ignoring parts of a model that were not traced). With an example of bacterial ribosome crystal structure, I show that the *checkMySequence* can successfully identify errors in both protein and nucleic acid components of complex structures, resulting from model tracing issues and errors in reference sequences that would be otherwise very difficult to identify. Owing to its performance and full automation, *checkMySequence* is applicable to the analysis of very large models. For example, validation of a complete cryo-EM structures of 80S ribosome at 3.4 Å resolution discussed in this work, with 48 chains and over 11,000 protein and nucleic acid residues takes less than six minutes on a laptop.

## DATA AVAILABILITY

The doubleHelix program source code and installation instructions are available at https://gitlab.com/gchojnowski/doublehelix. The doubleHelix sequence assignment algorithm has been added to the *checkMySequence* program to enable comprehensive validation of sequence assignment in the models of protein-nucleic acid complexes. The updated method is available at https://gitlab.com/gchojnowski/checkmysequence. Corrected models of mammalian and bacterial ribosome structures presented in this work are available at https://doi.org/10.5281/zenodo.7650444.

## ACKNOWLEDGEMENTS

I would like to thank all the microscopists who decided to deposit half-maps to EMDB which was of great help while developing *doubleHelix*. I would also like to thank Sean Eddy for his help with the configuration of INFERNAL suite. I am very grateful to Daniel J. Rigden and Katherine S. H. Beckham for critical reading of the manuscript and very helpful comments.

